# Emerging many-to-one weighted mapping in hippocampus-amygdala network underlies memory formation

**DOI:** 10.1101/2023.09.06.556568

**Authors:** Jun Liu, Arron F Hall, Dong V Wang

## Abstract

Memories are crucial for our daily lives, yet the network-level organizing principle that governs neural representations of our experiences remains to be determined. Employing dual-site electrophysiology recording in freely behaving mice, we discovered that hippocampal dorsal CA1 (dCA1) and basolateral amygdala (BLA) utilize distinct coding strategies to represent novel experiences. A small assembly of BLA neurons rapidly emerged during memory acquisition and remained active during subsequent consolidation, whereas the majority of dCA1 neurons engaged in the same processes. Machine learning decoding revealed that dCA1 population spikes predicted the BLA assembly firing rate. This suggests that most dCA1 neurons concurrently index an episodic event by rapidly establishing weighted communications with a specific BLA assembly, a process we call “many-to-one weighted mapping.” Furthermore, we demonstrated that closed-loop optoinhibition of BLA activity triggered by dCA1 ripples after new learning resulted in impaired memory. These findings highlight a new principle of hippocampus-amygdala communication underlying memory formation and provide new insights into how the brain creates and stores memories.

## Introduction

Converging evidence suggests that the formation of new memories involves a complex neural network of multiple brain regions, including the hippocampal dorsal CA1 (dCA1) and basolateral amygdala (BLA). How neuronal populations within these brain regions encode and exchange information is yet to be fully understood [1-4]. The formation of episodic memories, or memories of events, can be divided into two processes: memory acquisition and consolidation. Memory acquisition is rapid and can index episodic events almost instantaneously; however, these memories are often fragile and only become stabilized after a slow transformation process known as memory consolidation [5, 6]. This process critically involves sleep, with dCA1 ripples, fast oscillations (100–300 Hz) that occur predominantly during slow-wave sleep and immobility, playing a crucial role [5-7]. Despite this understanding, it is still unclear how the dCA1–BLA circuitry enables such rapid indexing and gradual consolidation of new memories.

Memories of events consist of three primary components: where, when, and what [8, 9]. Previous studies have demonstrated that the dCA1 encodes where and when by sequences [10, 11]. In contrast, no consensus has been reached regarding how dCA1 neurons encode specific events (what). Given the continuous unfolding of episodic events in our daily life, the encoding of these events becomes intricately intertwined with recall of prior relevant experiences. Consequently, dCA1 neurons become highly entangled in the process of encoding both ongoing and past event information [12], making it challenging to disentangle dCA1 spike patterns that specifically encode individual events.

Presently, there are two major hypotheses regarding how the hippocampus encodes episodic events (what). One prominent view holds that “(event) information is sparsely encoded in distributed ensembles of hippocampal neurons” [13], although there is limited evidence supporting this sparse-coding hypothesis by dCA1 neurons [13-15]. In fact, recent findings suggest the contrary, revealing that a substantial portion of dCA1 neurons (up to 50%), instead of a minority, are activated during fear memory procedures [16-19]. Moreover, it has been shown that up to 30% dCA1 neurons exhibit co-activation upon exposure to two distinct contexts [16], indicating a substantial overlap of dCA1 neurons in encoding different memories.

The other influential view suggests that the hippocampus functions to link or bind, rather than encode event information that is otherwise represented in a distributed neural network [20, 21]. This viewpoint, however, fails to address the precise mechanisms through which dCA1 activity effectively achieves such linking/binding, and it has not been directly tested experimentally. To address this knowledge gap, our study employs a different approach to deduce dCA1 encoding principles by computing activity correlations between dCA1 neurons and selective BLA assembles. This approach leverages recent findings that BLA neurons form distinct assembles to represent salient events [22-24].

## Results

### Emerging dCA1 ripple–BLA communication underlies memory consolidation

To investigate the encoding principles and communication dynamics of the dCA1 and BLA neuronal populations, we conducted dual-site *in vivo* recording of the dCA1 and BLA (up to 16 tetrodes per site; Figure 1A). All mice received a contextual fear memory procedure that consisted of pre-training sleep, training, post-training sleep, and fear recall test (Figure 1B). Neuronal spikes and local field potentials (LFPs) were recorded throughout the entire process, which enabled us to study dCA1–BLA neuronal ensemble dynamics at each memory stage, including acquisition, consolidation, and retrieval.

**Figure 1.**
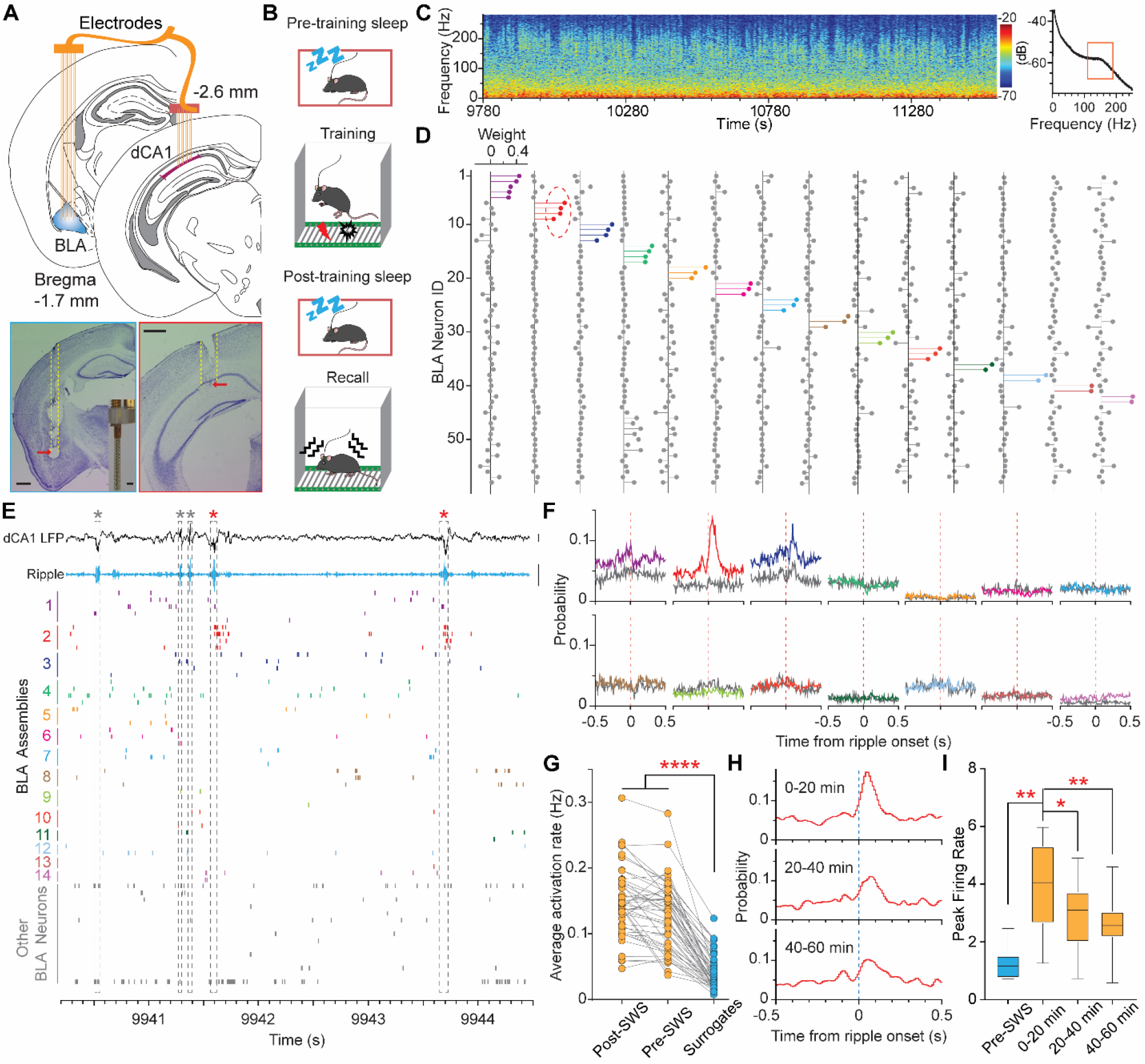
Emerging dCA1 ripple-to-BLA assembly communication during memory consolidation. **A**, Schematic of dual-site recording (top) and representative brain sections showing recording sites in the BLA and dCA1 (bottom). Inset, a self-constructed 16-tetrode array. **B**, Schematic of contextual fear memory procedure. **C**, Spectrogram (left), and power spectral density analysis (right) of representative dCA1 LFP recorded during post-training sleep. **D**, Independent component analysis identified 14 assembles (shown in colors) based on spikes of 58 BLA neurons recorded during the same sleep session as shown in C. **E**, Representative dCA1 LFP, band-pass filtered ripples (100–250 Hz), and spikes of the same 58 BLA neurons as shown in D. Note that assembly-2 exhibits robust activation after a subset of dCA1 ripples. **F**, Cross-correlograms between dCA1 ripples (n = 2147) and the 14 BLA assembles. Grey and color lines indicate pre- and post-training sleep, respectively. **G**, Activation rates of all identified BLA assembles (n = 48 from 5 mice) are significantly higher than chance during both pre- and post-training sleep. P < 0.0001, F_1.860, 87.42_ = 208.6, one-way ANOVA; ****P < 0.0001, Bonferroni post-hoc. **H**, Cross-correlograms between dCA1 ripples and BLA assembly-2 (as shown in E) across post-training 20-min sleep epochs. **I**, A gradual decay in the correlation between dCA1 ripples and BLA memory neurons (n = 10 from 5 mice) across the post-training sleep epochs. P = 0.0003, F_1.307, 11.77_ = 22.02, one-way ANOVA; **P = 0.0039, *P = 0.0108, **P = 0.0041, Bonferroni post-hoc.

We first identified slow-wave sleep (SWS) stages based on prominent dCA1 delta [25] and ripple oscillations (Figure 1C). Next, we conducted an independent component analysis (ICA) of population spikes recorded during SWS to identify major BLA assembles, i.e., small groups of co-activated neurons [26]. As an example, the ICA identified 14 assembles based on the activity of 58 BLA neurons recorded during post-training SWS (Figure 1D; Figure S1). We observed that one selective BLA assembly (#2), which we termed “BLA memory assembly,” robustly increased its activity after a subset of dCA1 ripples (Figure 1E). Subsequent cross-correlation analysis confirmed this observation and revealed additional BLA assembles that showed decreased (# 1&8), decreased followed by a rebound (#3), or little change of activity (Figure 1F). Notably, none of these BLA assembles showed dCA1 ripple-correlated activity during pre-training sleep (Figure 1F). This suggests the emergence of dCA1 ripple-to-BLA assembly communication underlying memory consolidation.

Next, we sought to determine if the BLA assembles preexisted prior to learning or if they were newly formed during the learning process. Our analyses revealed that nearly all BLA assembles were already highly active during the pre-training sleep (Figure 1G). This suggests that memory formation primarily recruits preconfigured or preexisting BLA assembles [3], albeit with a minor degree of membership swapping within certain assemblies. Additionally, we investigated the time course of dCA1-to-BLA communication and found that such communication was more prevalent during early-stage SWS (Figure 1 H&I).

### Memory-associated ripples are enlarged and elongated

We next asked if the dCA1 ripple-to-BLA assembly communication conveyed unique information. To address this, we classified dCA1 ripples into two categories: memory and non-memory associated ripples. Memory-associated ripples were defined if they occurred coinciding with BLA memory assembly activation, while the remaining ripples were defined as non-memory ripples (Figure 2A). Our analyses revealed that memory-associated ripples had significantly larger amplitude and longer duration (Figure 2B). Moreover, these memory-associated ripples contained distinct contents, i.e., spikes of distinct dCA1 neurons (Figure 2C). Comparing memory-associated ripples to non-memory ones, most dCA1 neurons showed further firing changes on top of their already increased activity: about one third of them increased activity while another one third decreased activity significantly (Figure 2D). These findings suggest that dCA1 sends specific information to a selective BLA assembly for memory consolidation.

**Figure 2.**
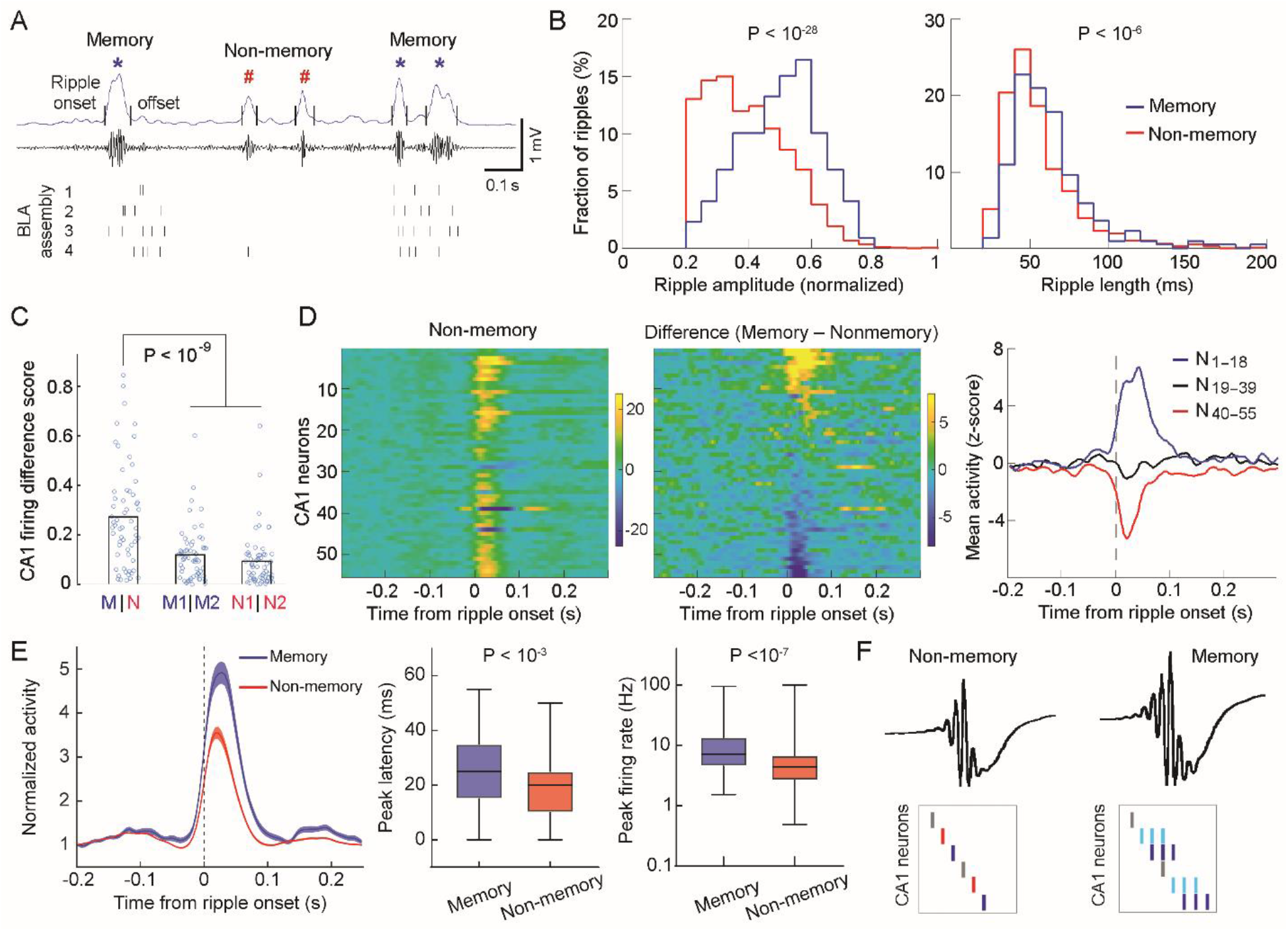
Memory-associated ripples are enlarged and elongated. **A**, The BLA assembly (assembly #2 as shown in Figure 1 D/E) exhibits activation at larger/longer ripples. Overall, 219 memory and 2390 non-memory ripples were classified in this post-training recording session (∼1h). **B**, Memory-associated ripples have significantly larger amplitude (left) and longer duration (right). **C**, Memory-associated ripples convey distinct contents. M/M1/M2 and N/N1/N2 indicate memory and non-memory ripples, respectively (see **Methods**). **D**, Left, z-scored activity of dCA1 neurons (n = 55; recorded simultaneously with the 58 BLA neurons as shown in Figure 1 D/E) in relation to the onset of non-memory ripples. Middle, activity difference of the same 55 dCA1 neurons in relation to memory *vs*. non-memory ripples. The neurons are arranged in the same order in the two heatmaps. Right, mean activity of the activated (#1–18), unmodulated (#19–39), and inhibited dCA1 neurons (#40–55). **E**, Left, Mean activity (± s.e.m.) of all dCA1 neurons that were activated during memory or non-memory ripples (n = 105 *vs*. 171 neurons, from 5 mice). Middle & Right, dCA1 neuronal activity was higher and peaked later at memory-associated ripples. **F**, Our proposed model of large/elongated ripples that contain distinct contents. Red/blue ticks denote dCA1 neurons that decrease or increase activity; light blue ticks denote dCA1 neurons that show emerged activity; grey ticks denote dCA1 neurons that show little activity change during memory-associated ripples. All statistics are Wilcoxon rank-sum test.

We observed that the increased dCA1 activity during memory-associated ripples tend to be prolonged (Figure 2D, right). To verify this, we conducted an unbiased analysis on all dCA1 neurons that increased activity during either memory or non-memory ripples. Our analyses confirmed that dCA1 neuronal firings were elongated and more robust during memory-associated ripples across animals (Figure 2E). These elongated dCA1 ripples and enhanced firings likely signify the consolidation of newly formed memories [27]. Taken together, we propose a model of ripple-associated memory consolidation, as depicted in Figure 2F. During non-memory ripples, most dCA1 neurons exhibit a baseline probability of activation that possibly represents preconfigured rigid motifs [28, 29]. Upon memory consolidation, relevant dCA1 neurons exhibit robust increase and/or prolongation of activity during ripples, while other dCA1 neurons show decrease or no change of activity, which collectively represent specific information coding.

### dCA1 population spikes predict BLA memory assembly firing rate

We next asked if dCA1 population spikes can predict the firing rate of BLA assembles or neurons. To address this, we implemented generalized linear model (GLM) machine learning decoding [30]. Given that dCA1 neuronal activity preceded BLA memory assembly activity by ∼35 ms (Figure 3 A&B), we used a shift of 35 ms or longer between dCA1 and BLA neurons for the GLM decoding. Overall, dCA1 and BLA neuronal spikes across multiple sliding windows (lasting 100 ms; Figure 3C) before or after dCA1 ripples were extracted: 50% of them were used for training of the GLM decoder, and the remaining 50% were used for decoding. We used correlation coefficients between the real and predicted firing rates to indicate prediction power, resulting in a scale between –1 and 1 (Figure 3D).

**Figure 3.**
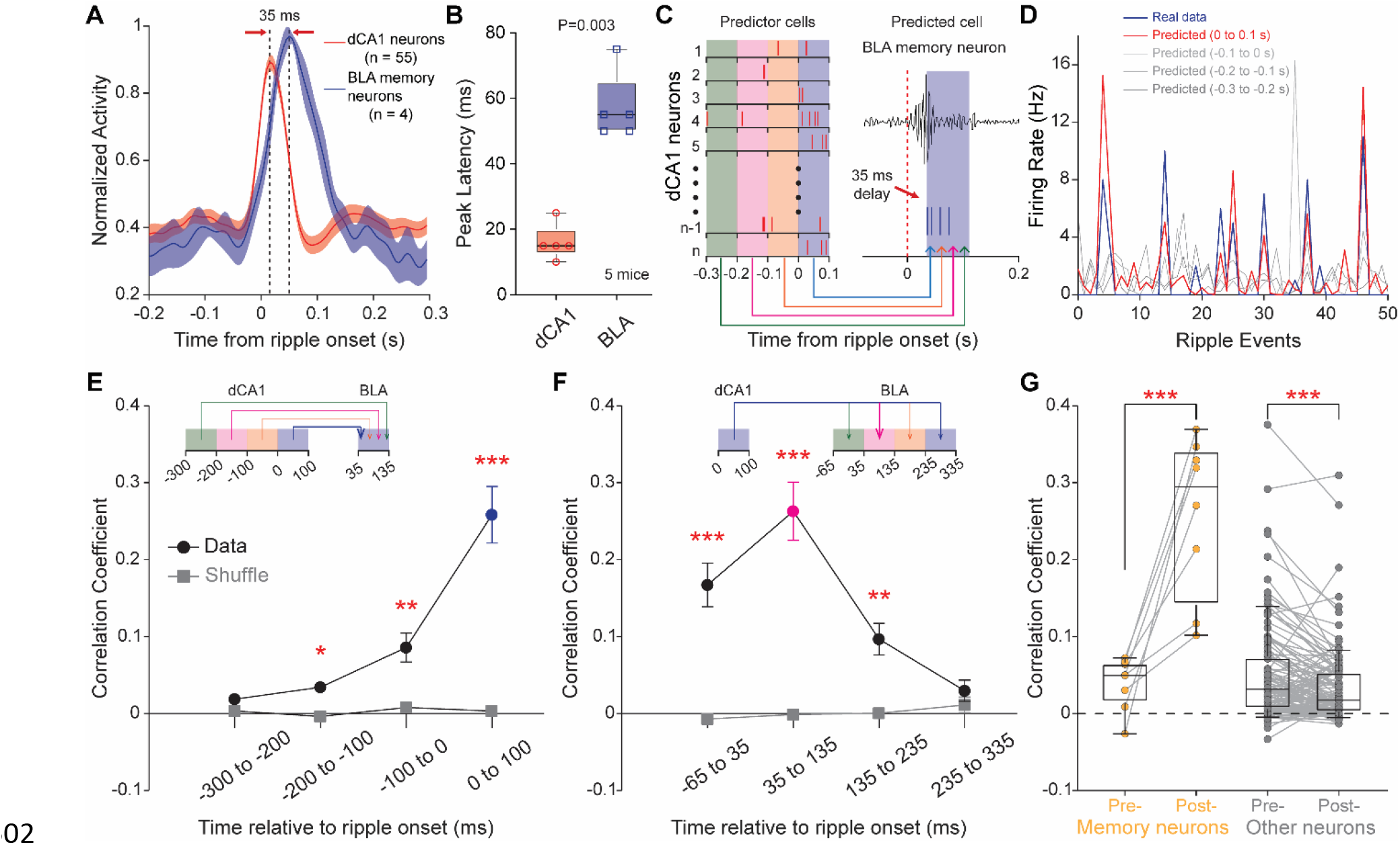
dCA1 population spikes can decode BLA assembly firing rate. **A**, Mean peri-ripple histograms (± s.e.m.) of simultaneously recorded dCA1 neurons and one BLA memory assembly. **B**, Peak firing rate latencies of dCA1 neurons and BLA memory neurons. **C**, Diagram of GLM decoding. Spikes of dCA1 and BLA neurons were binned at multiple 100-ms time windows. “Predictor cells” indicate dCA1 neurons; “Predicted cell” indicates a BLA assembly or neuron. **D**, Decoding the firing rate of one representative BLA memory assembly based on population dCA1 spike counts sampled at multiple bins as shown in C. **E&F**, Correlation coefficients between real and predicted firing rates of BLA memory neurons across multiple bins. Only recordings of 50 dCA1 neurons or more simultaneously (from 3 mice) were used for the decoding. Wilcoxon rank-sum tests revealed significant differences between the real and shuffled data at all time windows (P = 0.0148, 0.0030, 0.0002 for −200 to −100, −100 to 0, 0–100 ms windows in E; P = 0.0002, 0.0002, 0.0019 for −65 to 35, 35–135, 135–235 ms windows in F) except the -300 to -200-ms window in E (P = 0.1049) and 235–335 ms window in F (P = 0.6454). **G**, Correlation coefficients between real and predicted firing rates of BLA memory and non-memory (other) neurons during pre- or post-training sleep. ***P = 0.0008, ***P = 0.0002, paired *t* test.

Our findings demonstrated that the dCA1 population spikes predicted BLA memory assembly firing rate during the post-, but not pre-training sleep (Figure 3 D–F). This provides direct evidence of emerging weighted communications from dCA1 neurons to a selective BLA assembly in memory formation. In contrast, dCA1 population spikes had limited prediction on non-memory neurons, although the prediction power on a small subset of them appeared to be high during both pre- and post-training sleeps, which may reflect pre-existing dCA1-to-BLA communications (Figure 3G). Interestingly, the overall prediction power on non-memory neurons decreased after learning (Figure 3G, right), suggesting a bias towards the consolidation of newly acquired memories and skipping older ones, at least during the early-stage sleep.

### Emerging many-to-one weighted mapping underlies memory formation

Previous attempts to directly characterize dCA1 spike patterns that encode specific episodic events, such as receiving shocks or air puffs [2, 31], have not yielded a framework for understanding dCA1 information coding principles. Therefore, we employed a different approach to deduce the encoding principle of dCA1 neurons by analyzing their relationship with selective BLA assembles. This leverages our findings of robust BLA memory assembly activity during all memory stages that include acquisition, consolidation, and retrieval (Figure 1; Figure S2). Our results revealed that most dCA1 neurons exhibit BLA assembly-correlated activity during training and post-training sleep, but not pre-training sleep (Figure 4A). This suggests the involvement of a significant proportion of dCA1 neurons compared to a small assembly of BLA neurons in memory formation and consolidation.

**Figure 4.**
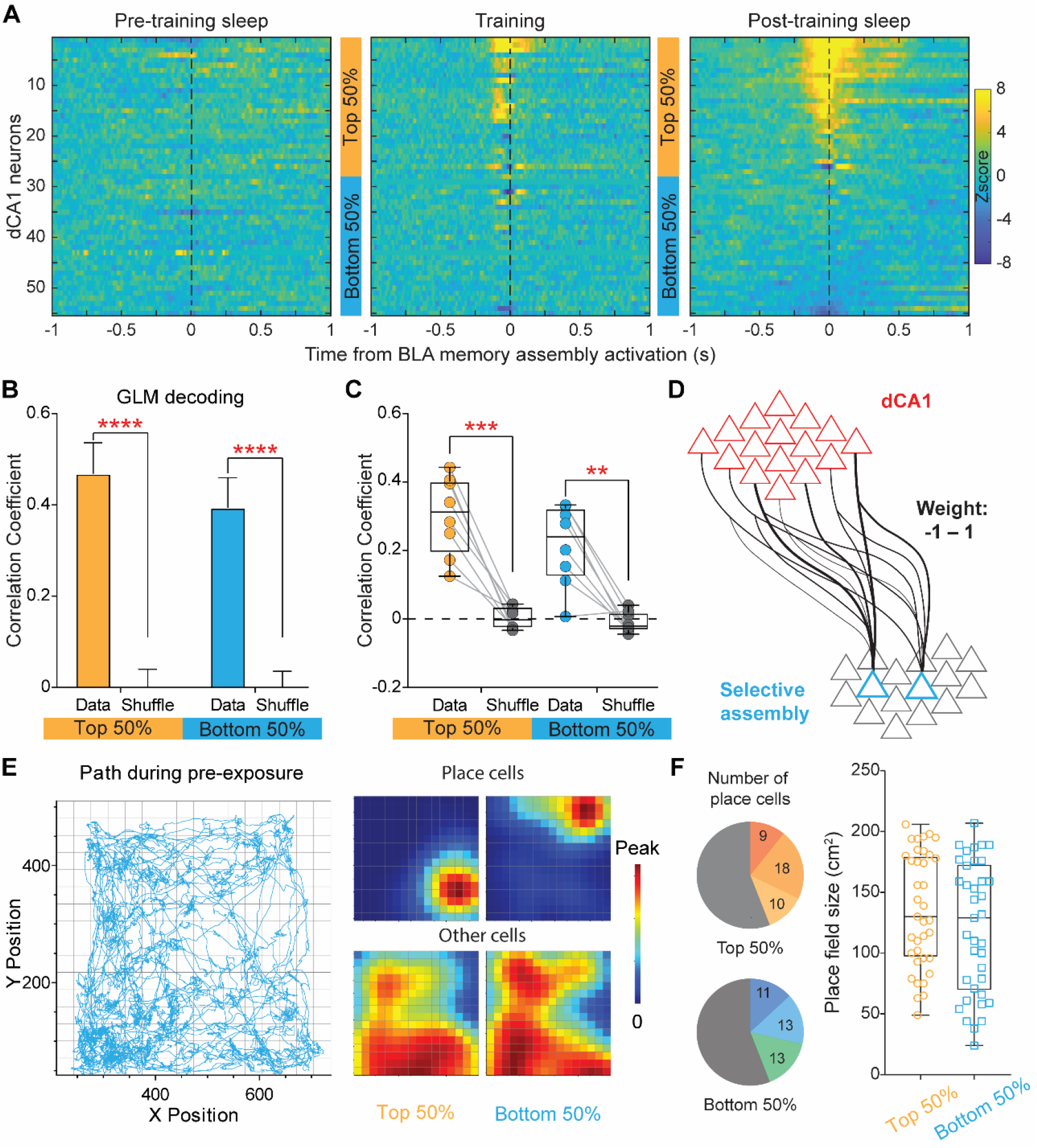
Emerging many-to-one weighted mapping underlies memory formation. **A**, Cross-correlogram heatmaps between one BLA memory assembly and simultaneously recorded dCA1 neurons during the pre-training sleep (left), training (middle), and post-training sleep (right). Neurons are arranged in the same order in the three heatmaps. **B**, Both the top 50% (corresponding to neurons #1–27 shown in A) and bottom 50% neurons (#29–55) predicted the firing rates of the BLA assembly, in comparison to the shuffled data (repeated for 100 times). **C**, Both the top 50% and bottom 50% neurons predicted the firing rates of individual BLA memory neurons (n = 8). Only recordings of 50 dCA1 neurons or more simultaneously (from 3 mice) were used for the decoding. **P < 0.01; ***P < 0.001; ****P < 0.0001; paired *t* test. **D**, Proposed model of many-to-one weighted mapping from dCA1 to BLA. **E**, A representative navigation path (left) and firing rate heatmaps of 4 example dCA1 neurons. **F**, Left, number of place cells among the top or bottom 50% dCA1 neurons from individual mice (n = 3; shown in different colors). Right, similar place field sizes between the top and bottom 50% place cells (P = 0.2180, unpaired *t* test).

Next, we divided the dCA1 neurons into subgroups based on their high or low correlated activity with the BLA memory assembly (Figure 4A). Our GLM decoding results showed that not only the higher-, but also lower-weight dCA1 neurons predicted the BLA assembly firing rate (Figure 4 B&C). This suggests that the decoding of the BLA assembly activity involves a contribution from many dCA1 neurons, rather than a few. Taken together, we propose a model of emerging many-to-one weighted mapping from dCA1 neurons to a selective BLA assembly that underlies the formation of a new memory (Figure 4D).

Additionally, we conducted place cell analysis. Our results showed that subsets of dCA1 neurons can function as place cells, with similar place field sizes between the higher- and lower-weight neurons (Figure 4 E&F). These results suggest that memory coding (non-spatial) and spatial coding are represented by partially overlapping groups of dCA1 neurons [32, 33].

### Closed-loop optoinhibition during post-training sleep impairs memory

To determine if dCA1 ripple-coincided BLA activity is necessary for memory consolidation, we employed a closed-loop optoinhibition approach. We microinjected AAV-CaMKII-stGtACR2 [34] into the BLA bilaterally and then implanted two optic fibers slightly above the injection sites, which enabled later optoinhibition of BLA pyramidal neurons. Meanwhile, we implanted four tetrodes into the dCA1 to record ripple activity (Figure 5A). After 2–3 weeks to allow viral expression, mice underwent a contextual fear conditioning procedure, followed by closed-loop, delayed, or no-stimulation of the BLA (Figure 5B). This procedure lasted two hours during post-training rest/sleep, similar to that reported previously [35, 36].

**Figure 5.**
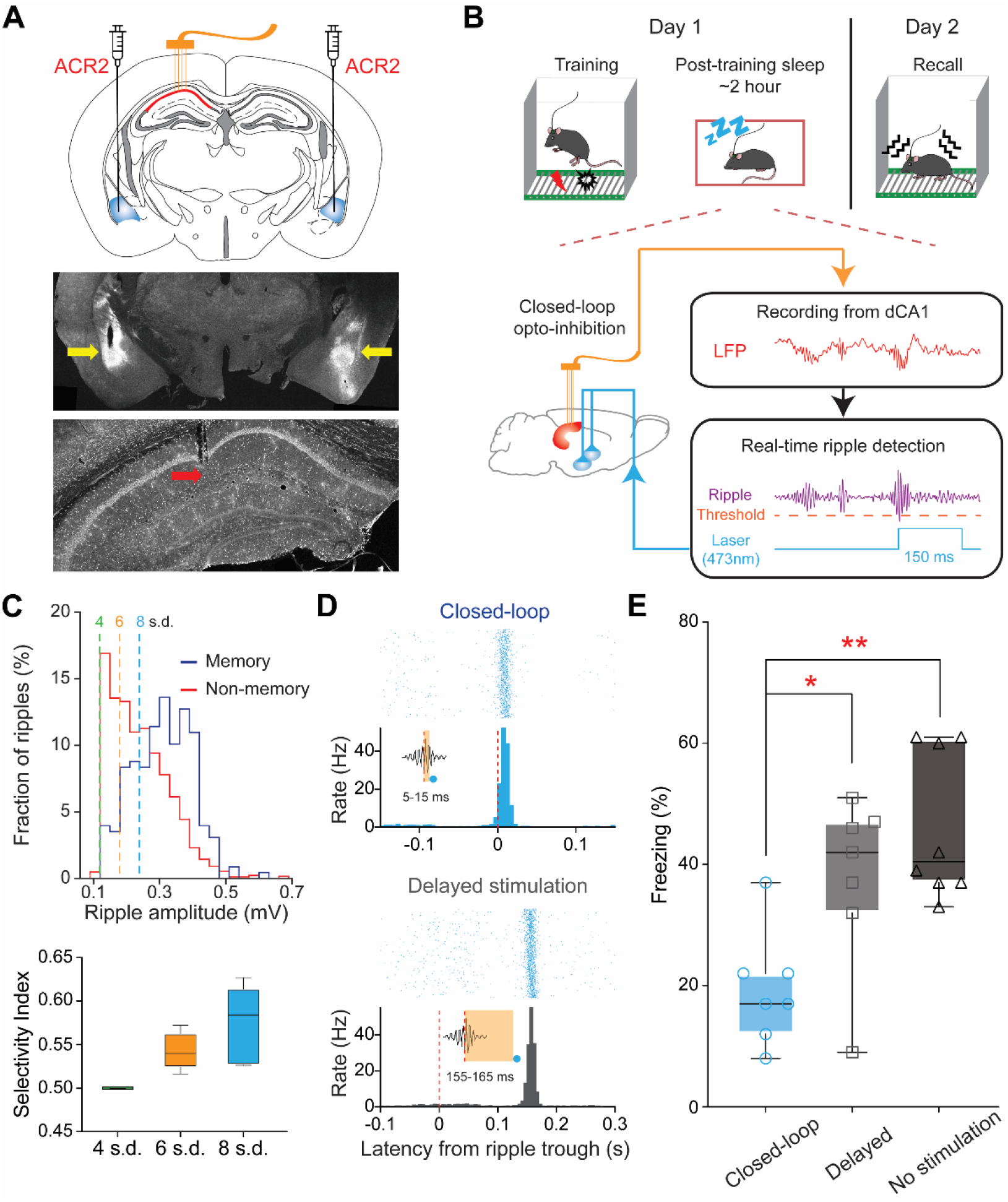
Closed-loop optoinhibition during sleep impairs memory consolidation. **A**, Schematic of surgical procedure (top), and representative coronal sections showing viral expression mainly in the BLA (middle) and recording site in the dCA1 (bottom). **B**, Schematic of contextual fear memory procedure and closed-loop optoinhibition. **C**, Ripples detected at higher threshold (8 *vs*. 6 *vs*. 4 s.d.) have higher selectivity of memory-associated ripples. Selectivity index = M/(M+N). M (or N) indicates the percentage of above-threshold ripples among memory-associated (or non-memory) ripples. **D**, Optoinhibition latencies of the closed-loop group mice (top) and delayed-stimulation group (bottom). **E**, Closed-loop group mice shows impaired fear memory compared to delayed- or no-stimulation groups (P = 0.0013, F_2, 19_ = 9.583, one-way ANOVA; *P = 0.0303, **P = 0.0011, Bonferroni post-hoc).

Our analysis showed that large-amplitude dCA1 ripples had a greater impact on memory consolidation (Figure 2B). Therefore, we used a high threshold (8 s.d.) to trigger closed-loop optoinhibitions, which preferentially targeted memory-associated ripples (Figure 5C). Our offline analysis confirmed that the closed-loop group mice received the optoinhibition within a short latency of ∼10 ms after ripple detection, whereas the delayed group had an extended latency of ∼160 ms (Figure 5D). Behaviorally, the closed-loop group mice exhibited significantly reduced freezing compared to the delayed or no-stimulation groups, indicating impaired contextual fear memory (Figure 5E).

## Discussion

Our findings suggest that most dCA1 neurons concurrently index an episodic event by rapidly establishing weighted communications with a specific BLA assembly. We refer to this process as “many-to-one weighted mapping.” The involvement of a significant portion of dCA1 neurons in encoding a specific event presents several theoretical advantages. Firstly, this mechanism offers immense encoding capacity due to the numerous potential combinations of dCA1 weighted mappings. Secondly, it provides robustness and redundancy, thereby enhancing single-trial learning and minimizing potential disruptions from background or external noises [9, 37]. It is noteworthy that single-trial learning and memory formation are recognized as features of biological intelligence in comparison to long-training sessions of artificial intelligence [38].

In essence, we propose that many-to-one weighted mappings are responsible for creating distinct memories in network representation. Specifically, when intermittent dCA1 firings occur that represent any significant mental or cognitive process spanning a few hundred milliseconds, they constitute an exclusive population activity pattern. This patterned firing enables the establishment of a unique weighted mapping from many dCA1 neurons to a specific BLA assembly that represents the relevant significant event, likely through rapid long-term potentiation [39]. In other words, distinct dCA1 population activity patterns communicate with corresponding BLA assembles to rapidly index various episodic memories. It is possible that one dCA1 population activity pattern simultaneously communicates with multiple neuronal assembles that are distributed across the brain through collateral projections [40]. Consequently, dCA1 reactivations during ripples bind these distributed assembles to form a network representation of long-term memories. In support, we have demonstrated that closed-loop optoinhibition of the BLA activity triggered by dCA1 ripples following a new learning experience impairs relevant memory.

Only a few studies have investigated hippocampus-amygdala interactions at the neuronal ensemble level during memory formation [2, 3]. One study found that dCA1 neurons increased correlation with selective BLA neurons during fear memory formation [2]. Our study extended this finding by demonstrating that population dCA1 spikes predicted the firing rate of a selective BLA assembly during ripples. Another study found that the BLA assembles are preconfigured prior to memory acquisition [3]. Our results confirmed this and showed that the activity of these preexisting BLA assembles was not driven by dCA1 ripples. We speculate that the recruitment of preexisting BLA assembles to establish communication with the dCA1 enables rapid and efficient circuit mapping. Together, an emerging hippocampus-to-amygdala communication appears to underline emotional memory acquisition and consolidation.

Anatomically, dCA1 and BLA are not directly connected by synapses [41, 42]. Accordingly, we found that the dCA1-to-BLA information flow took about 35 ms, which indicates a di-synaptic communication. It appears that only the entorhinal, perirhinal, and ectorhinal cortices receive direct inputs from the dCA1 and project directly to the BLA, based on comprehensive anterograde and retrograde tracing studies shown in Allen Brain Connectivity and Mouse Connectome Project (Figure S3). Theoretically, information relayed from the dCA1 to BLA can utilize a many→fewer→one *or* many→many→one principle (Figure S3). The former notion gains support in findings from immediate early gene studies, which have demonstrated that a large portion of dCA1 neurons (up to 50%) are activated during memory processing, whereas a moderate portion of entorhinal neurons (∼25%), and only a small portion of BLA neurons (∼10%) are activated in the same process [16-18]. These differences are further amplified with decreasing neuron numbers, from ∼400k in dCA1, ∼330k in deep-layer entorhinal cortex, to only ∼83k in BLA of rats [43-46].

Our research also demonstrated that memory-associated dCA1 ripples are elongated, which is consistent with a recent study showing that prolonging dCA1 ripples after learning improves memory [27]. Furthermore, we found that the elongation of ripples is accompanied by an overall prolongation and enhancement of neuronal firings in the dCA1, with some neurons increasing while others decreasing their activity. Additionally, our results revealed that memory-associated ripples were enlarged in amplitude, contrary to a previous study [27], which may reflect species differences between mice and rats [47]. Together, our findings suggest that elongated and enlarged ripples signify the consolidation of newly acquired memories, while shorter and smaller ripples may indicate baseline activity or preconfigured dCA1 rigid motifs [28].

## Methods

### Mice

Male C57BL/6 mice were purchased from the Jackson Laboratory (stock #000664). Mice were 3–4 months old at the time of surgery; after surgery, they were singly housed in cages (40 × 20 × 25 cm) containing corn cob and cotton material and kept on a 12 h light/dark cycle with *ad libitum* access to food and water. Experimental procedures were approved by the Institutional Animal Care and Use Committees of Drexel University and were in accordance with the National Research Council *Guide for the Care and Use of Laboratory Animals*.

### Stereotaxic surgery

Surgery procedures were similar to that used in our lab [23]. In brief, mice were anesthetized with ketamine/xylazine mixture (∼100/10 mg/kg, i.p.) and kept on a heating pad at 37°C. For *in vivo* electrophysiology recording, mice received implantation of two electrode arrays (8–16 tetrodes each) into the BLA and dCA1, respectively [48]. For closed-loop optoinhibition, mice received microinjection of AAV viruses (AAV1-CKIIa-stGtACR2-FusionRed; 0.25 μl; *Addgene* 105669) and implantation of two optic fibers (diameter 200 μm) into the BLA bilaterally; meanwhile, they received implantation of 4 tetrodes into the dCA1 unilaterally. The BLA coordinates were AP –1.7 mm, ML 3.4 mm, DV 3.9 mm; the dCA1 coordinates were AP – 2.6 mm, ML 1.8 mm, and DV 1.1 mm.

### *In vivo* electrophysiology

Each tetrode consisted of four wires (90% platinum 10% iridium; 18 μm diameter; *California Fine Wire*). A microdrive was used to couple with the electrode bundle, similar to that used in our lab [23, 25]. Neural signal was preamplified, digitized, and recorded using a *Blackrock* CerePlex or *Plexon* acquisition system; meanwhile, animals’ behaviors were recorded. With the *Blackrock* system, the local field potentials (LFPs) were digitized at 2 kHz and filtered at 500 Hz low cut; spikes were digitized at 30 kHz and filtered between 600–6000 Hz. For the *Plexon* system, the LFPs were digitized at 1 kHz and filtered between 0.7–300 Hz; spikes were digitized at 40 kHz and filtered between 400–7000 Hz. The tetrode arrays were gradually lowered daily until we recorded clear ripples and a substantial number of neurons; otherwise, mice were excluded from further analyses. The recorded spikes were sorted using the MClust 3.5 [23]; key dataset was manually verified using *Plexon* Offline Sorter. In total, spikes from five mice were used for analyses in this study; the neuron numbers in BLA and dCA1 were 58/55, 41/63, 36/52, 35/18, and 14/15, respectively.

### Assembly detection by independent component analysis (ICA)

ICA was performed like that described previously [26]. In brief, spike counts of each neuron were binned at 25 ms and z-scored to generate a neuronal population activity matrix (neurons × bins). Coactivity patterns were then extracted from this data matrix in two major steps. Firstly, the number of significant coactivation patterns was estimated by calculating the principal components (PCs) of the data matrix with variances above a threshold derived from an analytical probability function for uncorrelated data (Marchenko-Pastur distribution; Figure S1). Secondly, fastICA was used to extract the coactivity patterns from the projection of the data matrix into the subspace spanned by the significant PCs. In simple terms, this method first finds the significant PCs and then rotates them to match the ideal assembly patterns. These detected assembly patterns often were comprised of a small number of neurons with high weights, along with a larger group of neurons with low or zero weights. Neurons whose weights exceeded 2 s.d. from the mean weight of each assembly pattern were classified as members of an assembly (Figure 1D). The activation strength of each assembly was calculated by projecting the columns of the z-scored spike matrix onto the axis defined by the corresponding assembly pattern. The activation events were identified when the activation strength exceeded 5 s.d. from the mean activation strength of each assembly (Figure S1B). To investigate the significance of activation event rates, we computed the activation strength of surrogate assembles (500 surrogates for each assembly) generated by randomly permuting the data matrix (Figure 1G; Figure S1B). For all activation strength calculations, the assembly patterns extracted from post-training SWS sessions were utilized as templates.

### Ripples

Ripples were band-pass filtered between 100–250 Hz and ripple envelope was smoothed with a Gaussian kernel of 32 ms [49]. Ripple amplitudes were defined as the peak values of ripple envelopes (Figure 2A): those with amplitude exceeding 3 s.d. above the mean were used for analysis in Figure 2. In Figures 1/3/4, ripple amplitudes exceeding 5 s.d. above the mean were used for analysis. Ripple onsets and offsets were defined when ripple amplitudes exceed 1 s.d. above the mean before and after corresponding ripple peaks (Figure 2A). Ripple length was defined as the duration between the ripple onset and offset: only those longer than 20 ms were used for further analysis.

### Memory and non-memory associated ripples

We first identified BLA memory assembles or neurons if: 1) they showed significant activations within 150 ms after dCA1 ripple onsets during post-but not pre-training sleep (Figure 1F); and 2) they showed robust activation after receiving footshocks (>5-fold above baseline lasting 10 min or longer; Figure S2). Ripples (recorded during the post-training sleep) were defined as memory-associated ripples if the BLA memory assembly or neuron showed high activation (2 s.d. above the median) within 150 ms of the ripple onset. The remaining ripples were defined as non-memory ripples (Figure 2A).

**Ripple contents** (related to Figure 2C). M (or N) indicates the mean firing rate for each dCA1 neuron ±100 ms from memory-associated (or non-memory) ripples. Firing difference score M|N was defined as abs(M–N)/(M+N). M1|M2 was defined as abs(M1–M2)/(M1+M2), in which M1 and M2 indicate mean firing rates for each dCA1 neuron with randomly selected spikes (±100 ms from memory associated ripples). N1|N2 was similarly defined as abs(N1–N2)/(N1+N2).

### GLM decoding

We constructed generalized linear models (GLMs) with a log link function to predict spike counts of single BLA assembles or neurons during ripples based on population spike counts in dCA1 across specific time windows [30]. Ripples detected during pre- and post-training sleeps, and all dCA1 and BLA neurons were included for the analysis. Spike counts of each neuron or assembly were binned in 100-ms bins relative to ripple onset: −300 to −200 ms, −200 to −100 ms, −100 to 0 ms, 0 to 100 ms for dCA1, and −65 to 35 ms, 35 to 135 ms, 135 to 235 ms, 235 to 335 ms for BLA. We used dCA1 population spike counts in different time bins to predict the spike count of a single BLA assembly or neuron. We randomly partitioned the ripples into two equally sized datasets: one of them was used to train the GLM decoder, and the other was used for the test. For the test phase, the model derived from the training phase was applied to the dCA1 population spike data in the test set, yielding predictions for the predicted BLA spike counts across ripples. Lastly, we conducted correlation coefficient analysis between the predicted and real BLA spike counts to measure GLM decoding power on a scale from −1 to 1.

### Place cell analysis

To quantify place cells, the firing rate maps were generated in 1 × 1 cm spatial bins in NeuroExplorer and smoothed by a Gaussian filter (filter width, 5 bins) before being sent to MATLAB for further analyses. Low-activity dCA1 neurons with peak firing rate within any given spatial bin <1 Hz were excluded from further analysis. A place field was identified by counting the number of spatial bins where a neuron’s firing rate surpassed 50% of its peak firing rate. A neuron with combined place fields of less than one third of the total exploration area was considered a place cell.

### Fear conditioning (*in vivo* recording)

The fear-conditioning chamber used in the experiment was a square chamber measuring 25 × 25 × 32 cm, with a 36-bar shock grid floor (*Med Associates*). The behaviors of the mice were recorded using either the Blackrock Neurotech NeuroMotive or Plexon CinePlex video system. During training, the mice were first allowed to explore the footshock chamber for ∼3 minutes. They then received up to 10 mild footshocks (0.75 mA, 0.5 sec), with a 2–3 min interval between shocks. Note that we used direct-current footshocks to minimize electromagnetic noise, so occasionally mice missed a shock if they stood on two positively or negatively charged grids. One minute after the last shock, the mice were returned to their home cages. Approximately 2 hours later, the mice were placed back in the footshock chamber for a 5-minute contextual fear test. Neural activity was recorded continuously, including the pre-training sleep (1–2 hours), training (∼0.5 hour), post-training sleep (1–2 hours), and contextual fear test (5 min).

### Fear conditioning (closed-loop optoinhibition)

Three groups of mice were used: 1) closed-loop; 2) delayed; and 3) no-stimulation groups. All mice were singly housed and received daily handling for 1–2 weeks. All mice underwent a contextual fear procedure that consisted of three footshocks (0.75 mA, 2 sec; 1–1.5 min apart), similar to that described previously [50]. Freezing behaviors were automatically scored using *Med Associates* VideoFreeze [50]. To conduct closed-loop optoinhibition, we used the *Open Ephys* recording system and *Opto Engine* lasers. Only large dCA1 ripples (raw 100–250 Hz filtered traces) with peak amplitude exceeding 8 s.d. [51] were used to trigger bilateral optoinhibition of the BLA (0.5 mW; 150 ms), either immediately (closed-loop group) or after a delay of 150 ms (delayed-stimulation group).

### Histology

To mark the final recording sites, we made electrical lesions by passing 20-second, 10-μA currents through two tetrodes. Mice were deeply anesthetized and intracardially perfused with ice-cold PBS or saline, followed by 10% formalin. The brains were removed and postfixed in formalin for at least 24 hours. The brains were sliced into coronal sections of 50-μm thickness using *Leica* vibratome. Sections from the dual-site recording mice were stained with cresyl violet for microscopic examination of electrode placements, whereas other sections were mounted with Mowiol mounting medium mixed with DAPI for microscopic fluorescent examination of viral vector expression and optical fiber placements.

### Statistics

Sample sizes were based on previous similar studies in our labs [23, 25]. To determine firing-rate change during dCA1 ripples, the value that deviates from the mean by a *z*-score of >3.3 (P < 0.001) for at least three consecutive bins (bin = 5 ms) was considered significant. Other statistical analyses include Analysis of Variance (ANOVA) followed by post-hoc Bonferroni, Wilcoxon rank-sum test, and Student’s *t* test. All statistical tested are two-sided when applicable; P-values of 0.05 or lower were considered significant.

## Acknowledgements

We thank Dr. Vitor Lopes-dos-Santos for discussions on the ICA analysis and Drs. Gideon Rothchild and Hualou Liang for discussions on the GLM decoding. We thank Dr. Wen-Jun Gao for comments on an earlier version. This work was supported by the National Institutes of Health grants R01MH119102 (D.V.W.) and F31MH134582 (A.F.H.).

## Author contributions

Conceptualization & Methodology: LJ, DVW; Investigation: LJ, AFH, DVW; Writing – original draft: LJ, DVW; Writing – review & editing: LJ, AFH, DVW

## Data and materials availability

All data, code, and materials used in the analysis will be available upon publication of this manuscript.

## Supplementary Materials

Figs. S1 to S3

---

**Figure S1.**
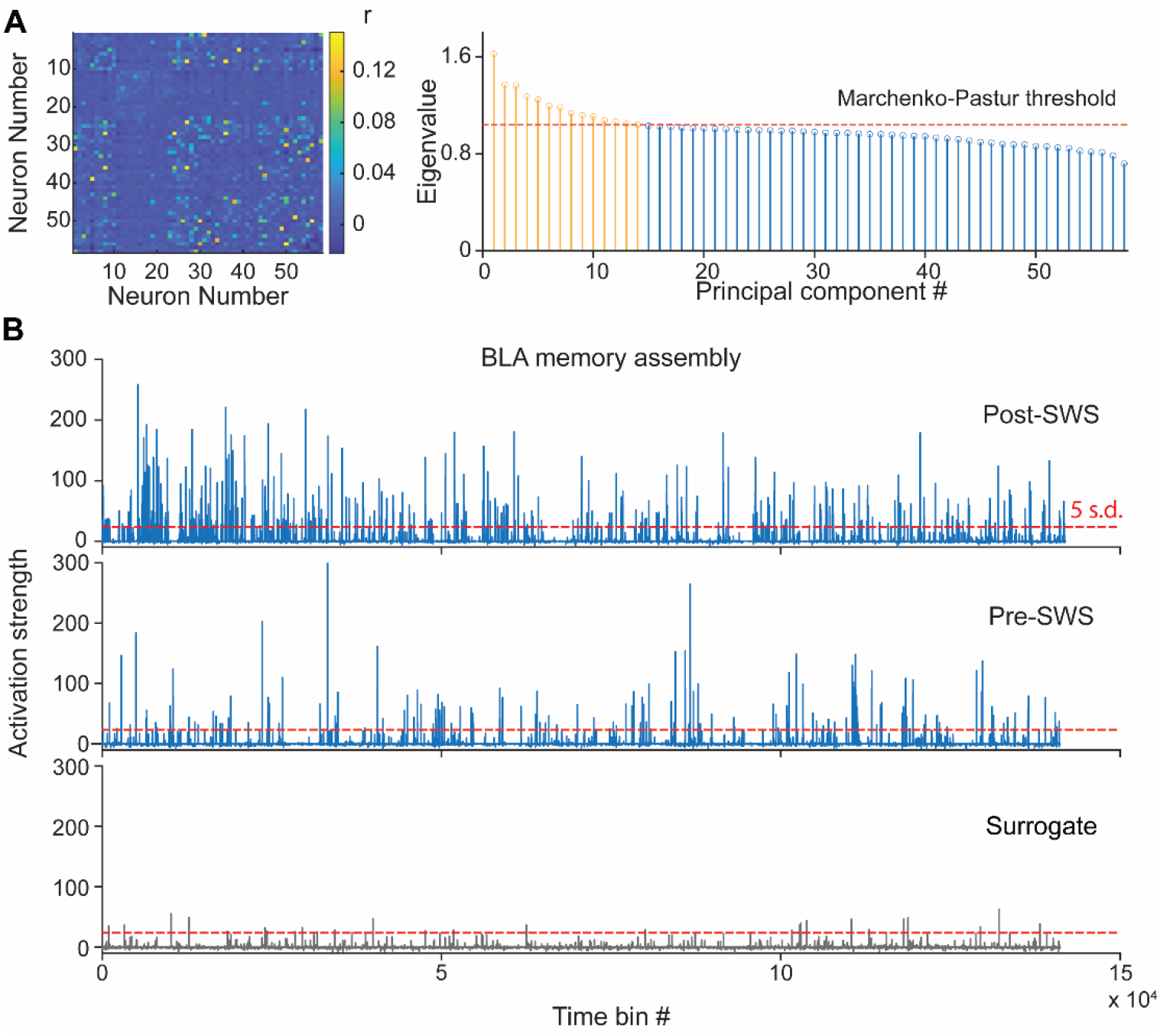
Independent component analysis (ICA). **A**, Correlation coefficient map of 58 BLA neurons (left; the same neurons as shown in Figure 1D) and Eigenvalues of the principle components (right; Marchenko-Pastur threshold was used to determine the number of assembles). **B**, Activity strength of assembly 2 (as shown in Figure 1 D/E) during pre- and post-training sleep and surrogate.

**Figure S2.**
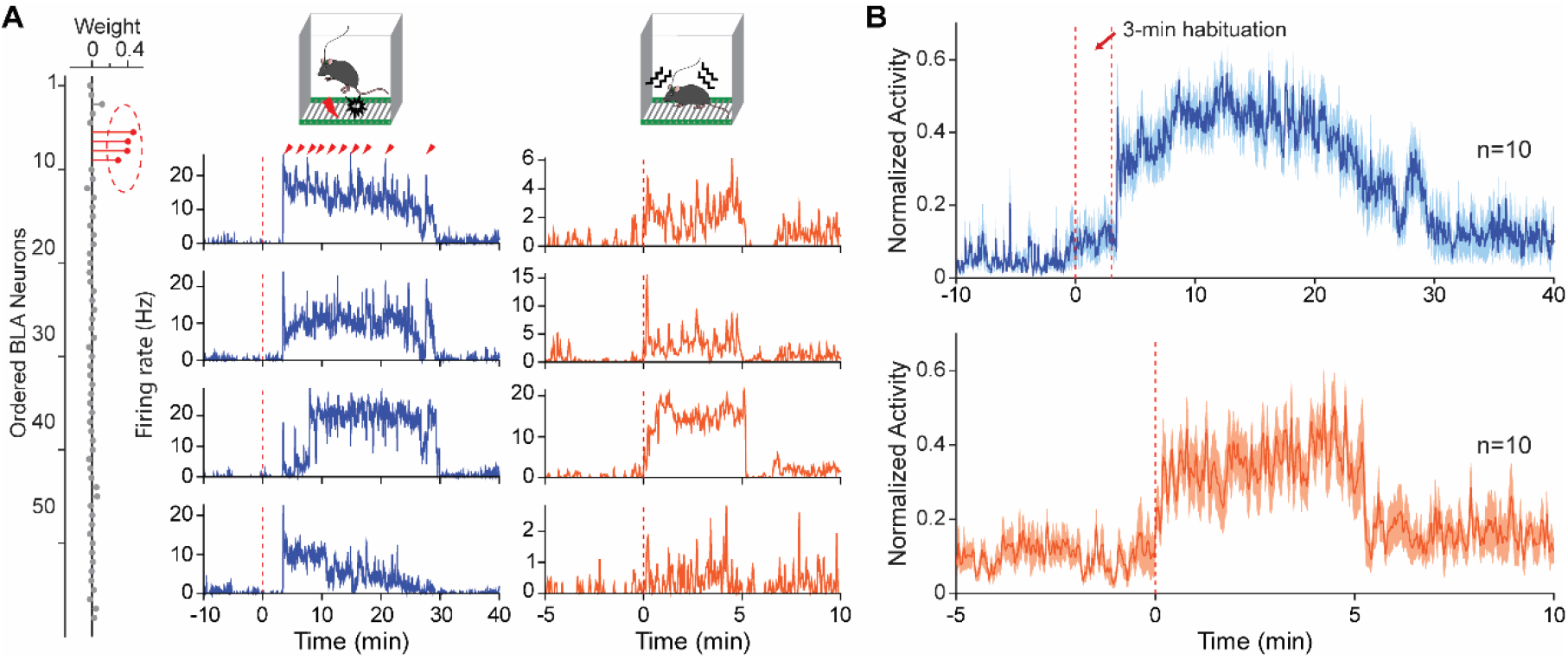
Emerged BLA memory assembly activity after footshock delivery. **A**, Left, the same assembly 2 (neurons #6–9) as shown in Figure 1 D/E. Middle & Right, firing rate histograms of the same 4 neurons during contextual fear training (middle; 10 footshocks) and memory retrieval (right). **B**, Mean activity (± s.e.m.) of BLA memory neurons during the training (top) and memory retrieval (bottom). Note that there’s little change of activity between 0–3 min before footshock delivery.

**Figure S3.**
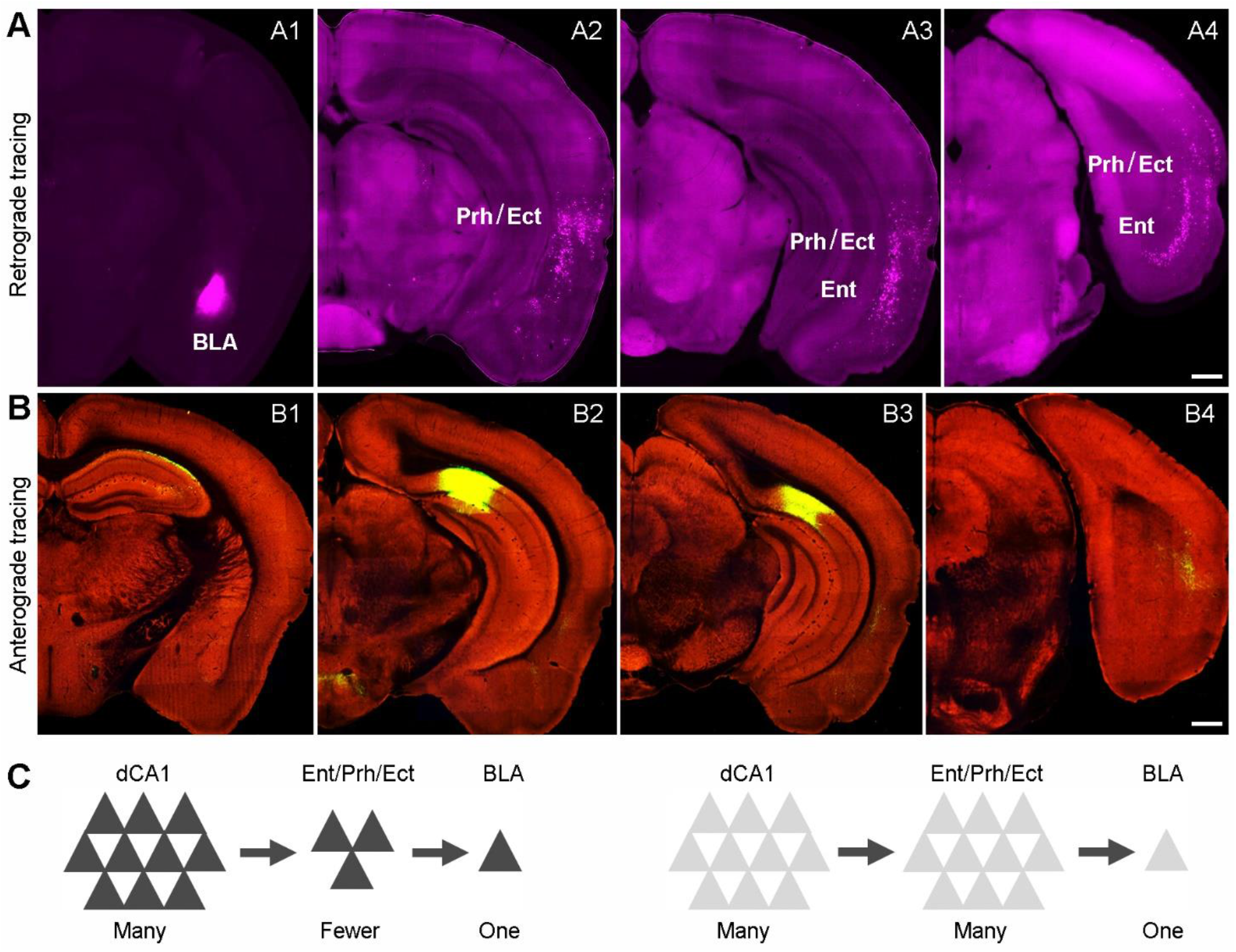
dCA1 communicates with BLA indirectly. **A**, Retrograde tracing of inputs to the BLA. Retrograde tracer CTB was microinjected into the BLA (A1). A2–A4, Retrograde labeling of neurons primarily in deep layers of entorhinal (Ent), perirhinal (Prh), and ectorhinal cortices (Ect). **B**, Anterograde tracing of dCA1 efferents. Anterograde tracer (AAV-Syn-EGFP) was microinjected into the dCA1 and adjacent subiculum (B2). Anterograde projection is seen mainly in the Ent (B3) and Prh/Ect (B2–B4), but minimal in the BLA. **C**, Two theoretical models of dCA1-to-BLA communication (many→fewer→one *vs*. many→many→one). Brain images are adapted from Mouse Connectome Project (A; https://cic.ini.usc.edu/) and Allen Brain Atlas (B; https://connectivity.brain-map.org/).

